# Emergence of a new material through ligation of DNA nanostar hydrogels

**DOI:** 10.64898/2025.12.21.695831

**Authors:** Audrey Cochard, Lucas Pinède, Yannick Tauran, Arnaud Brioude, Beomjoon Kim, Alexandre Baccouche, Vincent Salles, Anthony J. Genot, Soo Hyeon Kim

**Author notes:** Corresponding Authors: A. Cochard and S.H. Kim.

## Abstract

DNA nanostar hydrogels hold great promise as biomaterials, but their mechanical properties and nuclease susceptibility limit their use. Here, we propose ligation of nanostar sticky ends as a strategy to fundamentally transform the mechanical and dynamic properties of DNA nanostar hydrogels and enhance their nuclease resistance. Ligation eliminates free ends of DNA nanostars, conferring strong exonuclease resistance; suppresses strand rearrangement, converting liquid-like into gel-like assemblies; and prevents macroscopic reversibility despite preserving microscopic motif hybridization. Ligation also stiffens the material, enabling facile handling and 3D printing, and markedly slows digestion in serum-supplemented medium. In the presence of low concentrations of the DNase I inhibitor actin, ligated hydrogels remain stable for at least two weeks. As a proof-of-concept application, we captured and maintained cells within ligated hydrogels for one week in culture. By overcoming the limitations of DNA nanostar hydrogels, this work enables their use as programmable biomaterials that fully leverage the DNA nanotechnology toolbox.

## INTRODUCTION

Hydrogels are 3D hydrophilic polymer networks increasingly recognized as high-performance biomaterials for applications such as cell culture, tissue engineering and drug delivery^1^. They are broadly classified as natural or synthetic. Natural hydrogels, often inspired by the extracellular matrix, are inherently biocompatible and exhibit high bioactivity, but they suffer from batch-to-batch variability and limited tunability of their mechanical properties^1^. This is a critical drawback for cell culture or tissue engineering, where mimicking the material properties of specific tissues is crucial, as mechanical stimuli dictate cell functions^2–4^. By contrast, synthetic hydrogels, offer tunability but often lack biocompatibility and intrinsic bioactivity, and many cross-linking or functionalization strategies still rely on cytotoxic chemistries^5,6^.

Because conventional hydrogels rarely combine biocompatibility, tunability, and straightforward functionalization, DNA has emerged as an alternative material that addresses many of these limitations, owing to its well-defined structure, polymer properties, and the large toolbox developed to read, synthesize, and functionalize it. Initiated more than 40 years ago, DNA nanotechnology has become a mature field encompassing DNA nanostructure (e.g., DNA origami), DNA-based nanoparticle assemblies, and the more recent DNA hydrogels ^7–10^. Biocompatible, programmable and stimuli responsive, the latter have gained growing interest for biomedical applications, such as cell culture and capture, drug delivery, tissue engineering, biosensing and 3D bioprinting^11^.

DNA hydrogels can be either pure or hybrid, the latter incorporating DNA strands as bridges between non-DNA scaffold molecules, e.g. acrylamide chains^12^. Although pure DNA hydrogels can be more costly to synthesize, they offer finely tunable properties and ease of functionalization. For instance, gene-coding sequences can be ligated into the DNA network, or nanoparticles incorporated electrostatically, thereby enabling a wide range of applications, from protein- or RNA-producing gels to light-triggered drug release and shear-thinning hydrogels^13–16^. Moreover, sequence design can endow the material with specific responsiveness. For example, pH-sensitive DNA hydrogels have been engineered using interlocking sticky ends whose conformation varies with pH^17^. Similarly, incorporating both cell-specific aptamers and disulfide linkages, cleavable by the reducing agent GSH in the cytosol, into the design of DNA nanohydrogels has yielded systems for targeted gene regulation therapy^18^.

Multiple strategies exist to build DNA hydrogels. Rolling circle amplification (RCA) is a prominent approach in which the polymerization of long DNA chains leads to chain entanglement, optionally combined with hybridization^19–21^. Another attractive approach relies on base pairing between small DNA nanostar motifs that serve as building blocks, followed or not by ligation^22–24^. Base pairing is particularly appealing because of its simplicity: mixing as few as three complementary DNA strands bearing cohesive sticky-end (SE) sequences can trigger liquid–liquid phase separation (LLPS) and yield hydrogels above a concentration threshold. Since DNA hydrogel formation is governed by the sequence design, modifying the strands allows fine tuning of the hydrogel’s properties. For instance, lengthening the sticky-ends enhances both the mechanical strength and the sol–gel transition temperature, whereas introducing mismatched sites reduce them^24^.

Yet, two major challenges limit the broader application of DNA nanostar hydrogels: their susceptibility to nucleases, which are ubiquitous under physiological conditions, and their weak mechanical properties, which complicate handling and restrict their use, for example, in tissue repair^25^. These limitations originate from the SE-mediated connection between DNA nanostar motifs, which rely on reversible hydrogen bonds (energies of 1 - 3 kcal/mol versus approximately 100kcal/mol for carbon-hydrogen covalent bonds^26^). Manipulating the reversibility of these hydrogen bonds could therefore simultaneously enhance the mechanical strength of the hydrogel and improve its resistance to nuclease degradation.

Here, we propose sticky-end ligation as a novel strategy to fundamentally reprogram the mechanical properties, network dynamics, and nuclease resistance of DNA nanostar hydrogels. Systematic variation of the fraction of phosphorylated strands, i.e., strands competent for ligation, enables precise tuning of the extent of these transformations. We first found that ligation markedly enhanced the resistance of DNA hydrogels to exonuclease III by eliminating free DNA ends. Secondly, ligation converted reversible DNA droplets into irreversible materials. Thirdly, ligation prevented strand rearrangement, and ligated DNA hydrogels exhibited remarkable temporal stability and minimal fluorescence recovery after photobleaching (FRAP), both characteristic of a gel-like behavior which contrasts with non-ligated hydrogels liquid-like dynamics. Fourth, ligation substantially increased the mechanical robustness of the materials, enabling facile pipetting and compatibility with 3D printing. Finally, ligation conferred resistance to nucleases in serum-supplemented medium, which was further enhanced by addition of the DNase I inhibitor actin. As a proof of concept for potential applications, we demonstrated the suitability of ligated DNA hydrogels as three-dimensional scaffolds for cell culture, relevant for tissue engineering and artificial tumor reconstruction.

## RESULTS

### Ligation of DNA nanostar hydrogels

Each Y-motif strand carries an identical six-nucleotide palindromic sticky-end (SE) at the 5′ terminus, mediating interactions between Y-motifs and driving hydrogel formation (Fig. 1A). In this study, phosphorylation of the 5′ ends enabled ligation between adjacent Y-motifs upon addition of a ligase, generating new covalent phosphodiester bonds (Fig. 1B).

The three strands were mixed at 5 µM each in a 96-well plate and incubated for one hour at 37 °C with shaking at 450 rpm, yielding one hydrogel per well (700–1000 µm diameter). By varying the proportion of 5′-phosphorylated strands (from 0% to 100% of each Y-strand species), we generated hydrogels with different degrees of ligation upon ligase treatment (Fig. 1C and 1D, top). The resulting materials ranged from liquid-like, capable of rearrangement, fusion, and relaxation, to gel-like, with arrested dynamics (see infra). For consistency with commonly used terminology, we refer to all DNA-based materials as ‘hydrogels’ regardless of their material behavior. Hereafter, they are denoted according to their phosphorylation percentage, from P0 gels (with no phosphorylation) to P100 gels (all strands phosphorylated).

We evaluated ligation efficiency by monitoring degradation by exonuclease III (ExoIII). As foreseen, ExoIII fully digested P0 gels within one hour (Fig. 1D and 1E, left). Non-ligated P100 gels behaved similarly, indicating that phosphorylation alone did not confer enzymatic resistance (Fig. 1D and S1A). In contrast, ligated P20, P50, P80 and P100 gels persisted after treatment, with higher phosphorylation ratios correlating with reduced size loss, though ligated P100 gels still shrank by ∼50% (Fig. 1D and E). Similar trends were observed with X-motif hydrogels (Fig. S1B), though digestion proceeded more slowly, which may reflect a higher DNA density and/or greater network connectivity (Fig. S1C and D).

**Figure 1:**
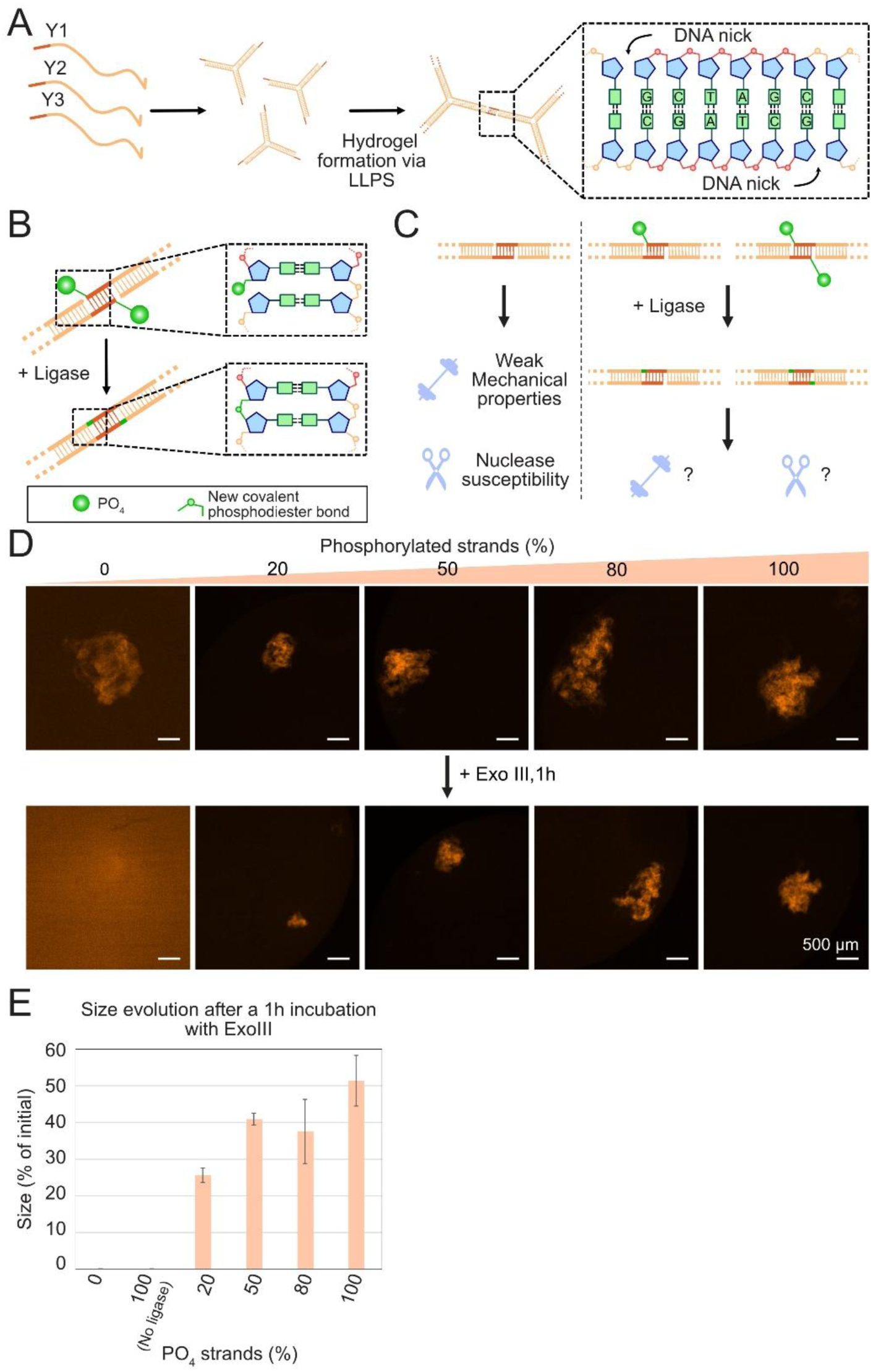
Formation and ligation of DNA droplets. **A.** Schematic of the formation of a DNA nanostar hydrogel through liquid-liquid phase separation. **B.** Formation of covalent phosphodiester bonds between Y-motifs via ligation. **C.** Comparison of phosphorylated-strands proportion on DNA hydrogels properties. **D.** Response of DNA hydrogels with different ratios of phosphorylated strands, after ligation, to one hour treatment with ExoIII exonuclease. **D.** Size evolution of DNA hydrogels following one hour treatment with ExoIII. From left to right, N = 6, 2, 3, 2, 2, 6.

If Y-motifs were fully interconnected and ligation was complete, full resistance to exonuclease digestion would be expected. Interestingly, hydrogels with higher phosphorylation ratios retained their overall shape (Fig. 1D, 80% and 100%), arguing against digestion initiated solely from free DNA ends at the surface. Instead, these results suggest homogeneous degradation by ExoIII within the hydrogel interior. The observed shrinkage may reflect limited penetration of the ligase—producing a ligated shell surrounding a non-ligated core—and/or an intrinsic connection degree lower than 100%, leaving some sticky ends unpaired and thus susceptible to exonuclease attack. However, if ligation were severely hindered in the core, more extensive internal degradation would be expected. Therefore, incomplete interconnection of Y-motifs appears to be the most plausible explanation for the size reduction.

### Ligation precludes DNA hydrogel reversibility

Y-motif–based DNA hydrogels assemble reversibly: heating the hydrogels over the sticky-ends melting temperature (T_m,SE_) disperse the motifs, and cooling allows for new SE bonding and formation of hydrogels (Fig. 2A, top)^23^. However, being over T_m,SE_ should not have any impact on fully ligated hydrogels, whose integrity does not rely anymore on SE bonding. As expected, while heating P0 gels over T_m,SE_ led to Y-motifs dispersion and a diffuse signal, P100 gels kept their initial structure (Fig. 2A). However, microscopic images of the DNA hydrogel immediately after reaching 100 °C revealed a disperse signal (Fig. 2B). Interestingly, upon cooling, PO gels re-formed, but P100 gels did not, indicating a loss of macroscopic reversibility (Fig. 2B).

To quantify thermal behavior, hydrogels formed at 37 °C were subjected to three melt–cool cycles (20→100°C, then 100→20°C, step of 0.5°C/2min). EvaGreen dye, labelling double stranded DNA, tracked hybridization dynamics.

All gels exhibited decreases in fluorescence at high temperatures (Fig. 2C, S2A and S2C). While the decrease was quite sharp for P0 gels and P100 gels, at about 70 and 84°C respectively for Y-motifs and 70 and 82°C for X-motifs, it spanned on a larger temperature range for partially ligated gels. We plotted the derivative of the melting curves (Fig. 2D and S2D). As expected, the curves of non-ligated Y-motifs based DNA hydrogels, P0 gels and non-ligated P100 gels, displayed a single peak centered around 70°C, the melting temperature of the Y-motifs as already highlighted in the literature (Fig. 2D and S2B). Interestingly, while 5’ phosphorylation did not influence the melting temperature of non-ligated Y-motifs based DNA hydrogels (Fig. S2B), it slightly decreased the T_m_ of the X-motifs (Fig. S2D), likely reflecting the more constrained structure of X-motifs hydrogels that could be slightly destabilized by the phosphate groups.

Ligation shifted the melting peak of fully ligated P100 gels ∼14 °C higher, consistent with longer ligated DNA structures. Partially ligated gels displayed two melting transitions. A first peak, centered around 70 °C, decreased with increasing phosphorylation and likely stemmed from non-ligated SE. Above 50% phosphorylation, this peak disappeared, as better revealed by the heatmap of the derivatives curves, indicating reduction to nil of non-ligated SE (Fig. 2E and S2E). A higher-temperature peak progressively appeared as from very low percentages of phosphorylated strands, at a temperature gradually increasing from around 78°C to 84°C (Fig. 2D, 2E). This peak likely arose from the ligated portion of the DNA hydrogels and may encompass both the fully and half ligated SE, explaining the gradual increase in temperature with more and more SE being fully ligated.

**Figure 2:**
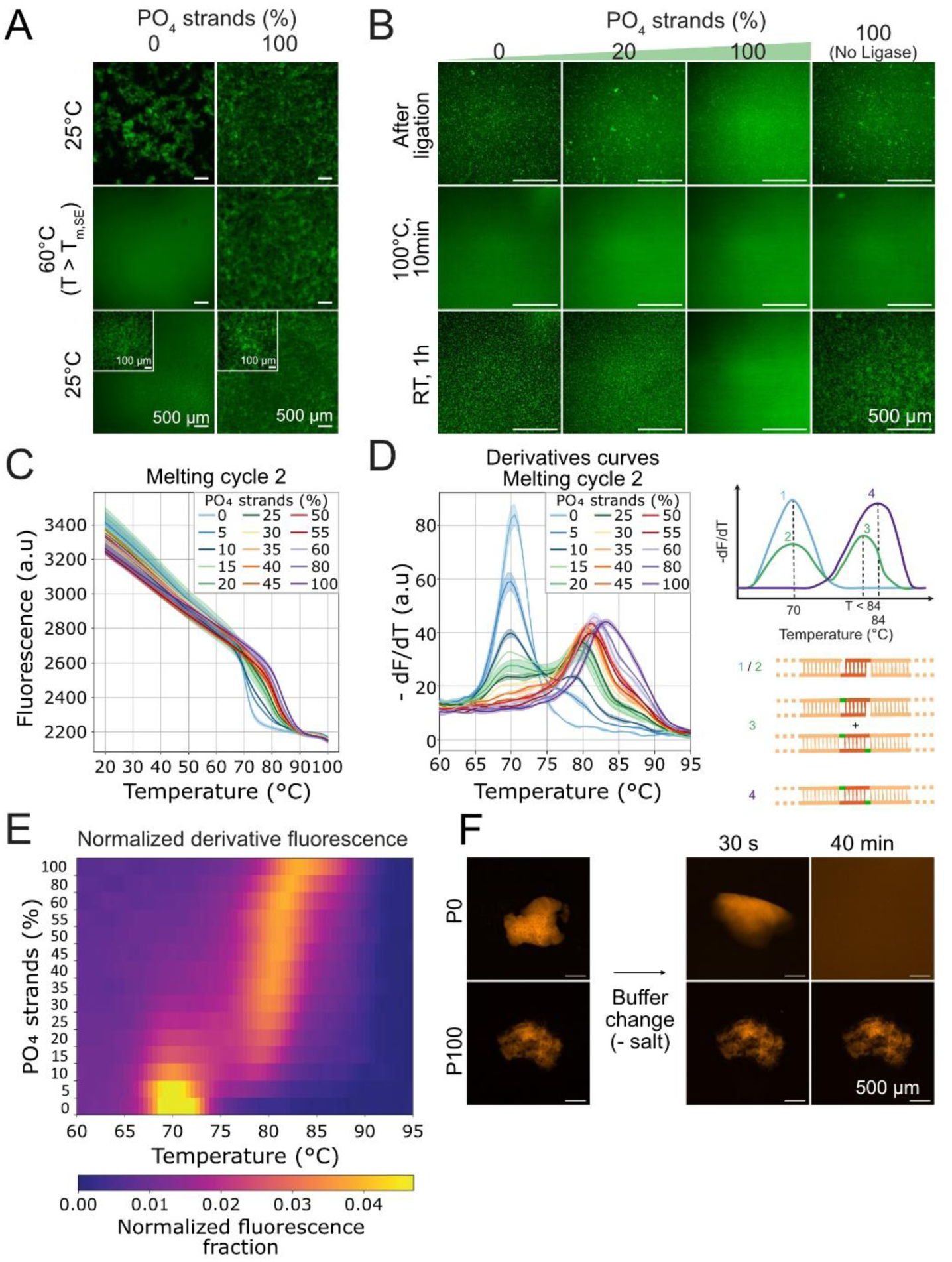
Loss of apparent reversibility. **A.** Microscope imaging of P0 and P100 gels while heated from RT to 60°C and back to RT. **B.** Microscope imaging of P0, P20, P100 and non-ligated P100 gels at RT, soon after being heated at 100°C for 10 min, and back at RT for 1h. **C.** Melting curves of the 2^nd^ cycle for P0, P20, P100 and non-ligated P100 gels. **D.** Derivative curves between 60 and 95°C of the melting curves of the 2^nd^ cycle displayed in C (left) and schematics of the melting events corresponding to each peak (right). **E.** Normalized derivative fluorescence intensity of the curves displayed in D. For each condition, derivative curves were baseline-corrected and normalized such that the total area under the curve equals 1. The color therefore represents the fractional contribution of each temperature point to the overall derivative fluorescence signal of that condition. **F.** Microscopy images of P0 and P100 gels (top and down, respectively), before and after washing the buffer for an unsalted buffer.

Despite the microscopic irreversibility of P100 gels, melt–cool cycling revealed consistent fluorescence jumps in every cycle, demonstrating reversibility at the level of hybridization, but not of macroscopic structure. A plausible explanation is that Y-motif networks contain interpenetrating loops, as described very recently in the literature^27^. Ligation might have frozen these topological constraints, so that heating denatured the strands but could not disentangle the network. Cooling may have allowed for the rehybridization of these interlocked structures. In support of this, replacing the buffer with an unsalted buffer completely dissolved P0 gels, while ligated P100 gels remained intact (Fig. 2F).

**Figure 3:**
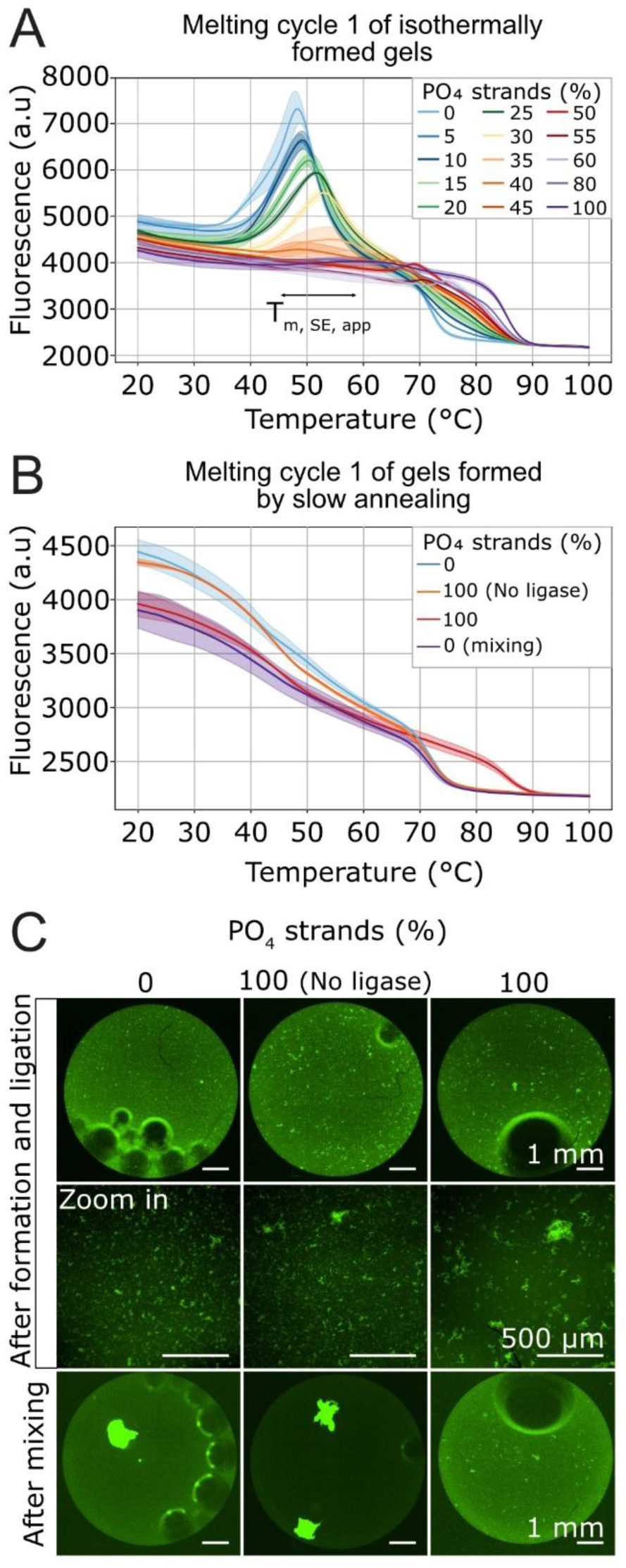
Influence of the formation method on the thermodynamical state. **A.** Melting curves of the 1st cycle (directly after formation and ligation) for P0, P20, P100 and non-ligated P100 gels. **B.** 1^st^ melting curves of DNA hydrogels formed by slow annealing. **C.** Microscopy imaging of P0, non-ligated P100 and P100 gels (from left to right) after formation and after mixing with a thermomixer for 1h. The second line is a zoom in of the first line.

### Melting the DNA hydrogels reveals the influence of the formation process over their thermodynamical state

In addition to high-temperature fluorescence drop, the 1^st^ melting curves exhibited another striking feature: a peak shifting from around 48°C for P0 gels to about 55°C for P35 gels, and disappearing at higher ligation levels (Fig. 3A). Fully phosphorylated but non-ligated gels displayed the same peak (Fig. S3A) as well as X-motif hydrogels (Fig. S3B). We attributed this peak to structural rearrangement within the hydrogels. Mixing the strands may have triggered a rapid gelation process, yielding kinetically trapped, random architectures rather than thermodynamically optimized ones. Increasing temperature would then promote transient SE unbinding/rebinding and relaxation into lower-energy configurations. Above T_m,SE_, SE unbinding domination would lead to the progressive dissociation of DNA motifs and fluorescence decrease. Partial ligation of sticky ends likely increased the effective T_m,SE_, accounting for the upward shift in peak position, and the peak ultimately disappeared when the network becomes too crosslinked to reorganize. Consistent with this interpretation, subsequent melting curves after slow cooling (Fig. 2C and S2A) or from hydrogels assembled by slow annealing (90°C to 20°C, -1°C/min, Fig. 3B), showed no low-temperature peak, indicating that annealed structures are already near equilibrium.

Microscopic images of DNA hydrogels further revealed that formation in PCR tubes yielded dispersed droplets rather than the single large hydrogels obtained in well plates (Fig. 3C, top). We thus examined whether slow-annealed hydrogels could be mixed post-formation to assemble into a single thermostable structure. Mixing led to the coalescence of non-ligated, liquid-like DNA material but not of the ligated, solid-like one (Fig. 3C). Importantly, the large non-ligated hydrogels formed after mixing displayed melting curves without the peak around 50 °C (Fig. 3B), demonstrating that a two-step process (slow annealing followed by mixing) produces a single, thermodynamically stabilized hydrogel.

### Ligation precludes Y-motifs rearrangement around the sticky-end melting temperature

DNA nanostar hydrogels display liquid to solid-like properties depending on SE length, motif valency and temperature. Longer SEs or higher branching numbers increase the temperature at which motif mobility decreases due to enhanced SE stability and network rigidity^23^. Here, the progressive disappearance of the peak around the SE melting temperature (T_m,SE_) in the melting curves (Fig. 3A) suggested that SE ligation reduces motif mobility near T_m,SE_.

Imaging DNA hydrogels over time at 37°C (T < T_m,SE_) and 45°C (T ≈ T_m,SE_) revealed a visible shape modification of P0 and P20 gels, with the rough surface obtained after mixing gradually smoothing out, driven by surface tension minimization (Fig. 4A). This process was faster at 45 °C (Fig. 4B), consistent with previous reports showing that these hydrogels behave as liquids at 45 °C but gels at 37 °C^23^. In contrast, gels with over 50% of phosphorylated strands retained their shape and size for at least four days (Fig. 4A and S4A), indicating that above a certain ligation threshold, SE detachment and subsequent rearrangement are effectively suppressed.

To probe motif mobility more directly, we performed widefield LED Fluorescence Recovery After Photobleaching (FRAP). Because the hydrogels studied here (_∼_800 µm diameter) were too large for localized bleaching, we used smaller hydrogel beads (_∼_50 µm) and monitored fluorescence recovery of a bleached half at 45 °C. As bleaching was performed with widefield LED illumination, recovery curves reflect relative mobility rather than absolute diffusion coefficients. As expected from macroscopic observations, P0 gels showed continuous fluorescence recovery over time, and did not reach a plateau after 20 minutes (Fig. 4C and D). Non-ligated P100 gels behaved similarly (Fig. 4D). In contrast, ligation drastically reduced recovery by _∼_40% in P20 gels, _∼_50% in P50 gels and P80 gels, and _∼_75% in P100 gels (Fig. 4D). For hydrogels with over 50% of phosphorylated strands, recovery reached a plateau within 5 minutes, further indicating a static, crosslinked structure.

The small size of the hydrogel beads also enabled fusion assays at 45 °C. P0 gels fused and relaxed within 3 min, typical of a liquid-like state (Fig. 4E). P20 gels still fused, but required about 20 min (Fig. 4F). Hydrogels with over 50% phosphorylated strands showed no fusion or relaxation (Fig. 4G). Together, these observations (macroscopic reshaping, FRAP, and fusion) demonstrate that ligation effectively suppresses SE dynamics and provides fine control over the mechanical and dynamical properties of DNA nanostar hydrogels.

**Figure 4:**
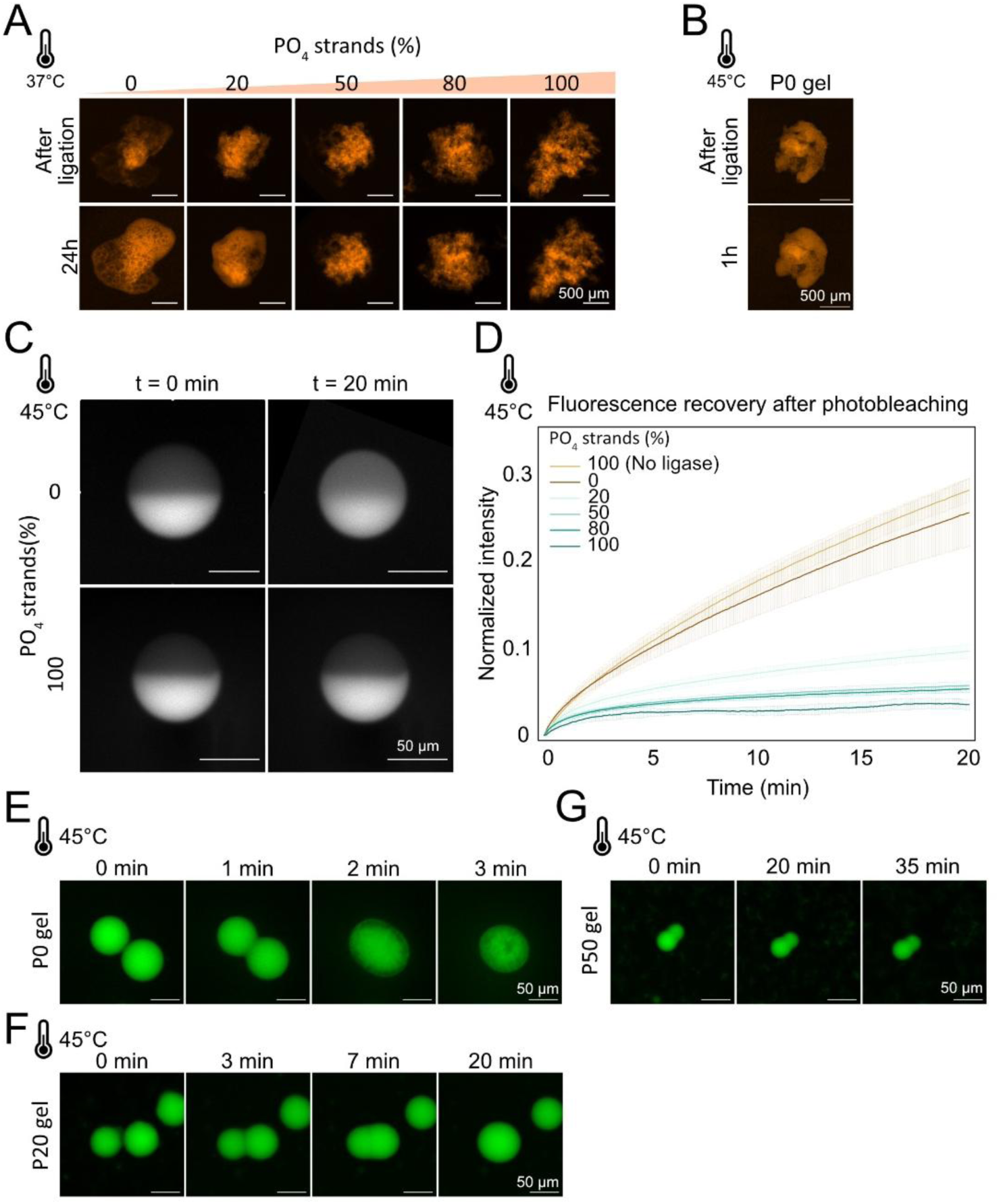
Thermodynamical behaviors of DNA hydrogels around T_m, SE_. **A.** Evolution at 37°C for 24h of DNA hydrogels with different percentages of phosphorylated strands, after ligation. Some images were rotated to better highlight the absence of DNA hydrogels shape evolution over time. **B.** Evolution of a P0 gel at 45°C for one hour. **C.** Representative image sequences of FRAP experiments carried out at 45°C on a P0 and a P100 gel (upper and lower images, respectively). **D.** Time series of the fluorescence intensity of DNA hydrogels after bleaching. **E. to G.** Representative sequential images of coalescence events for P0 (E), P20 (F) and P50 (G) DNA hydrogels at 47°C.

### Enhanced mechanical resistance of ligated DNA hydrogels

We next investigated whether the frozen topology of ligated DNA hydrogels affected their mechanical resistance. P0 and P100 gels were subjected to increasing shaking speeds in a thermomixer, from 500 to 1000 rpm. Non-ligated hydrogels remained stable up to 700 rpm, then gradually deformed, first rounding and then elongating at 800 rpm, and ultimately fragmented around 900 – 1000 rpm (Fig. 5A). The maximal shear stress in a rotating system can be estimated as τ_ω_ = *a*√(ρμω^3^), where *a* is the radius of rotation (m), ρ (kg.m^-3^) and μ (kg.m^-1^.s^-1^) are the density and dynamic viscosity of the medium, respectively, and ω is the angular velocity (rad.s^-1^)^28^. From this calculation, non-ligated hydrogels began to deform under shear stresses of approximately 2–2.5 Pa and shattered around 3–3.5 Pa (Fig. 5C). In contrast, P100 gels were unaffected by shaking and retained their overall shape even at 2000 rpm, the upper limit of our instrument (Fig. 5B, C, D).

**Figure 5:**
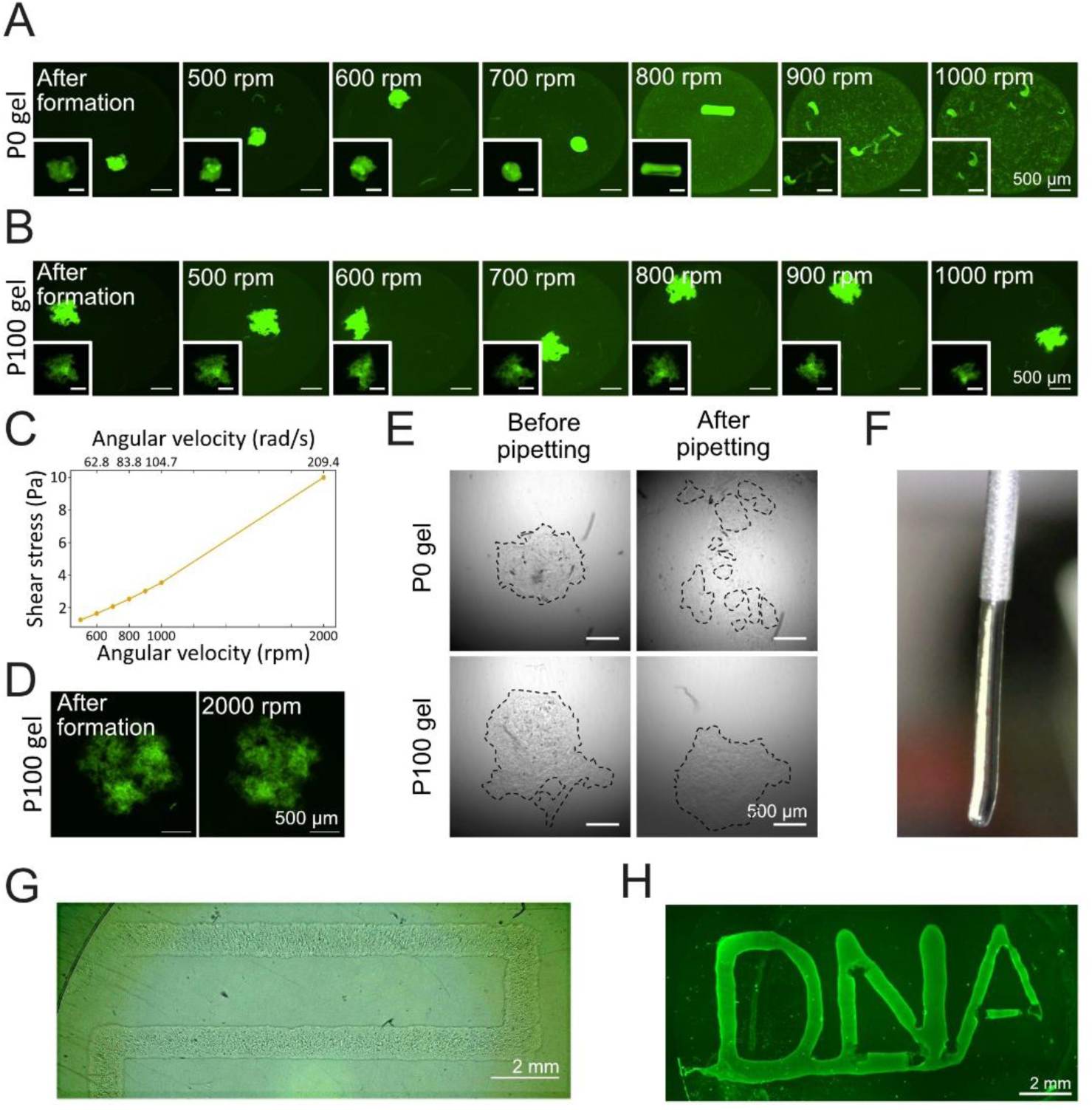
DNA hydrogels resistance to shear stress allows for their manipulation and 3D printing. **A** and **B.** Response of P0 (A) and P100 (B) DNA hydrogels to increasing speeds of rotation, from 500 to 1000 rpm. Hydrogels were submitted to each speed for 30 minutes, before increasing to the next speed. Whole field images are saturated so that fragments of DNA hydrogels in the well are visible. Inserts on the bottom left of each image correspond to unsaturated DNA hydrogels. **C.** Equivalence between angular velocity (rpm and rad/s) and shear stress (Pa). **D.** P100 DNA hydrogel submitted to rotation speeds up to 2000 rpm. **E.** Non-ligated and ligated DNA hydrogels (up and down, respectively) before and after undergoing two up and downs pipetting (left and right, respectively). **F.** Extrusion of a ligated DNA hydrogel through a G21 needle. **G.** Printing of a ligated DNA hydrogel with a G21 needle (internal diameter 0.514 mm) on a surface. **H.** Printing of a DNA hydrogel with a small nozzle (internal diameter of 0.1 mm).

This high mechanical resistance provides clear advantages for downstream applications. For instance, while P0 gels could not withstand pipetting, whose shear stress is of the same order of magnitude as the shaking assay^29^, preventing facile manipulation, P100 gels resisted pipetting with remarkable robustness (Fig. 5E). As a-proof-of-concept for potential applications in tissue engineering or artificial tumor reconstruction, we explored their extrusion-based additive manufacturing suitability. When extruded through a 21G needle (internal diameter 0.514 mm) without a receiving surface, P100 gels formed continuous, unbroken threads (Fig. 5F). They could also be deposited along predefined paths on poly-lysine treated coverslips and retained their shapes after immersion in medium (Fig. 5G). Using a smaller conical nozzle (inner diameter 0.1 mm) further improved printing resolution and yielded precise patterns (Fig. 5H). The ability to print DNA hydrogels with predefined designs opens possibilities in tissue engineering and regenerative medicine, where cells could be embedded within the hydrogel prior to printing.

### Endonuclease resistance of DNA hydrogels in serum-supplemented medium

The ubiquitous presence of nucleases in physiological environments is a hurdle for the use of DNA hydrogels in biological applications. Serum-supplemented medium contains many nucleases, notably the non-sequence-specific DNase I. Expectedly, P0 gels underwent almost complete degradation in this medium in 2 days (Fig. 6A and B). Although preheating at 75°C efficiently inactivated nuclease activity, allowing DNA hydrogels to retain their size over 4 days (Fig. S5A), this method is detrimental to cell culture^30^.

In contrast, P100 gels exhibited no substantial degradation within 4 days, before being slowly degraded (Fig. 6A and B). The mechanical resistance of ligated DNA hydrogels may account for this extended resistance by preventing structural distortion required for DNase I binding.

We next tested the effect of actin, a known DNase I inhibitor^31^. Non-ligated hydrogels were still fully degraded within a few days, even at high actin concentrations (Fig. 6C and D). Conversely, P100 gels remained intact for at least 14 days with no size reduction, even at low actin concentrations (Fig. 6E and F). Therefore, actin further improve ligated DNA hydrogel resistance to DNase I.

Because controlled enzymatic digestion may be desirable in some applications (cell release, etc.), we next induced degradation of ligated DNA hydrogels by exposure to high nuclease activity. We found that one hour of treatment with highly concentrated DNase I was sufficient to digest all DNA hydrogels (Fig. 6G), demonstrating that enzymatic breakdown remains accessible when required.

### Cell culture in ligated DNA hydrogels

By overcoming nuclease sensitivity, ligated DNA hydrogels open new possibilities for designing DNA-based cell culture matrices. As a proof-of-concept, we next assessed their ability to capture and culture cells in three dimensions.

To encapsulate cells within the DNA hydrogels, we applied a method recently developed by our laboratory^32^. Cells were first labeled with a biotinylated antibody and then conjugated to a Streptavidin-DNA complex displaying three SE moieties identical to those of the hydrogel-forming DNA strands (Fig. 7A). This approach efficiently captures more than 70% of the introduced cells^32^.

**Figure 6:**
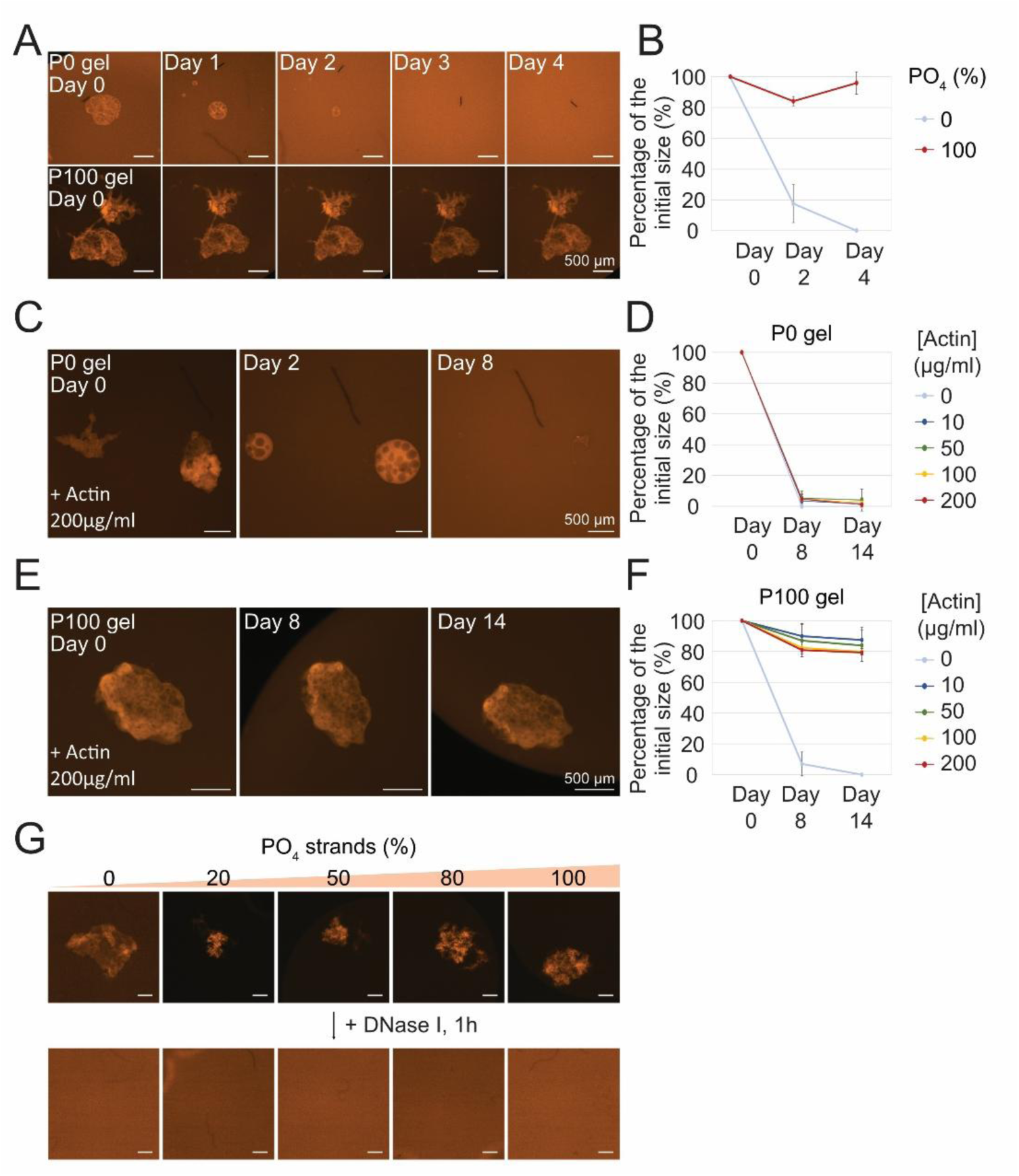
Evolution of DNA hydrogels in cell culture medium supplemented with FBS. **A.** Evolution of a P0 (top) and a P100 (down) gels in serum-supplemented medium over 4 days. **B.** Size evolution of P0 and P100 gels in serum-supplemented medium over 4 days. **C.** Evolution of a P0 DNA hydrogel in serum-supplemented medium with 200µg/ml of actin over 8 days. **D.** Size evolution of P0 hydrogels in cell culture medium supplemented with actin (concentrations ranging from 0 to 200 µg/mL). **E.** Evolution of a P100 gel in serum-supplemented medium with 200µg/ml of actin over 14 days. **F.** Size evolution of P100 hydrogels in cell culture medium supplemented with actin (concentrations ranging from 0 to 200 µg/mL). For each condition in **D.** and **F.**, N = 4. **G.** Response of P0, P20, P50, P80 and P100 gels, after ligation, to one hour treatment with DNaseI -XT endonuclease.

To monitor cell growth within the hydrogels, uncaptured cells must be removed to prevent late attachment to the hydrogel surface. Simple medium replacement was insufficient, with many cells remaining attached to the well bottom. We therefore leveraged the high mechanical robustness of P100 DNA hydrogels to pipetting and transferred the entire well content into a centrifuge tube, pelleted the hydrogels, discarded the supernatant, and resuspended the pellet in fresh buffer. Two washing steps were sufficient to eliminate almost all unattached cells, as confirmed by microscopic images (Fig. 7B).

We exploited the absence of strand rearrangement and the strong nuclease resistance of ligated hydrogels in serum-containing medium supplemented with actin. Cells remained within the hydrogel and viable for at least one week (Fig. 7C), and division of cells was observed, confirming that the hydrogels support cell viability (Fig. 7C, inset). Cell proliferation was relatively slow, even at higher initial seeding densities.

Because cell–matrix interactions are key regulators of proliferation, incorporating bioactive ligands could improve cell behavior. As a proof-of-concept demonstrating the modularity of our system, we functionalized P100 hydrogels with the RGD peptide, the most common integrin-binding motif in extracellular matrix proteins. We conjugated the streptavidin–DNA complex (displaying three SE) to biotinylated RGD peptides and confirmed efficient incorporation into the hydrogel network (Fig. 7D). Notably, the RGD peptide accumulated around captured cells (Fig. 7D, inserts), consistent with binding to cells surface integrins. Although we did not assess the biological effects of this modification here, these results illustrate the ease with which bioactive cues can be introduced, paving the way for future studies on cell–matrix interactions in DNA hydrogels.

**Figure 7:**
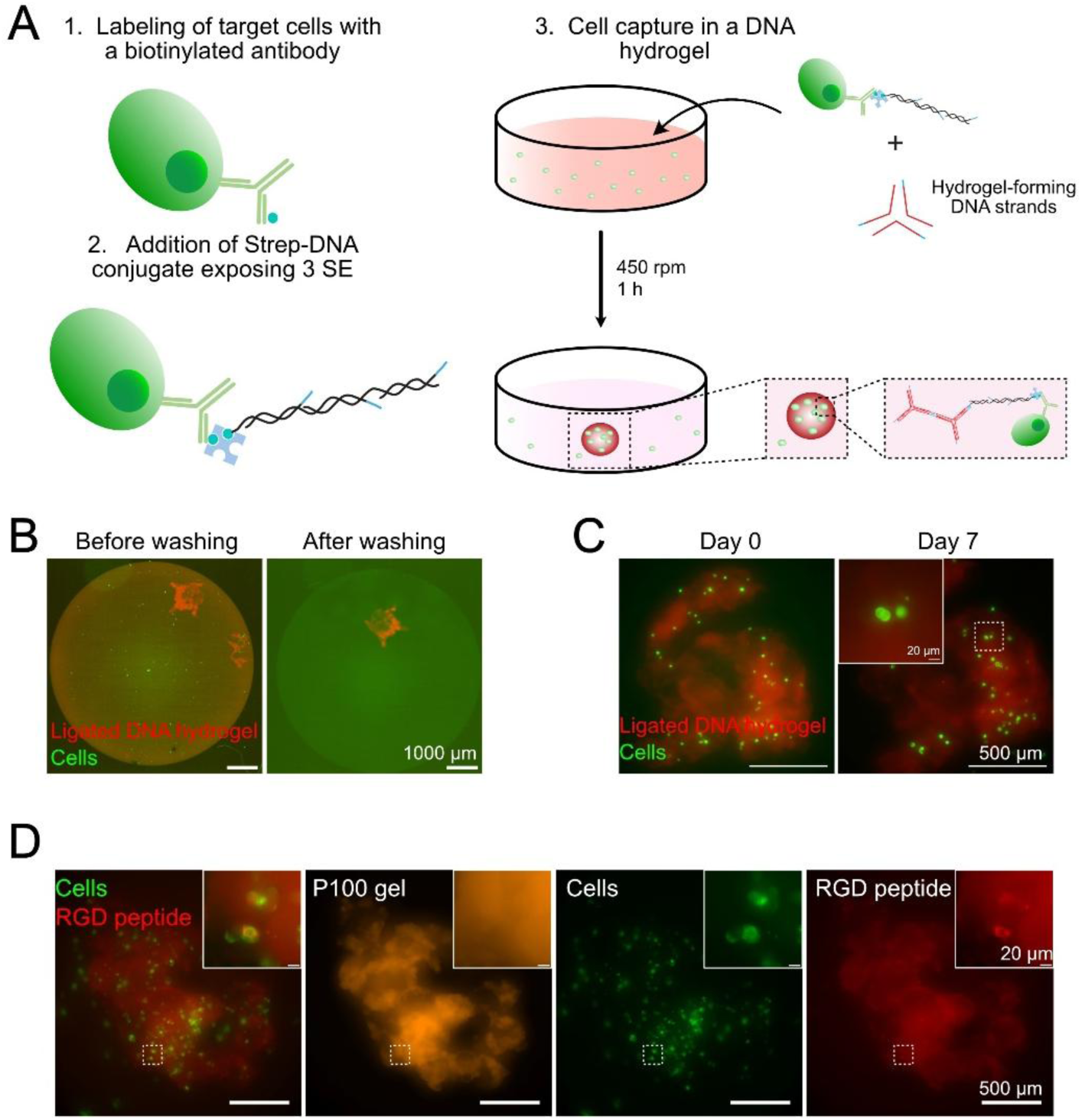
Cell culture in DNA hydrogels. **A.** Method for cell capture. Adapted from Bourdon et al. (2025). **B.** Representative epifluorescence images of a ligated hydrogel (red) and unbound cells in the supernatant (green) before and after washing by pipetting and centrifugation (left and right, respectively) **C.** Representative images of MCF-7 cells (green) captured in a P100 DNA hydrogel (red), on the capture day (left) and 7 days after (right). Insert: Zoom on a cell division event. D. Fluorescence imaging of MCF-7 cells (green) captured in a P100 gel (orange) functionalized with RGD peptides (red). Inserts are zoom in of the region delineated by a dashed white rectangle.

## DISCUSSION

DNA hydrogels are promising biocompatible materials for their self-assembly properties, tunability and ease of functionalization. Here, we demonstrate that ligation of DNA nanostar hydrogels profoundly alters their physical and biological properties. Ligation suppresses motif rearrangement, enhances mechanical robustness, and confers resistance to nucleases, ultimately enabling their use as stable scaffolds for 3D cell culture.

Those changes might stem from the transition from a dynamic and reversible network, maintained solely by SE hybridization, to a covalently crosslinked structure. By freezing the topology, ligation eliminates the SE binding-unbinding events, accounting for both the reduced molecular mobility, and the enhanced mechanical resistance. The creation of new covalent bonds also removed many DNA free ends, conferring a high resistance to exonucleases. Resistance was not complete, which may reflect partial connectivity between nanostar motifs. Interestingly, ligation led to an apparent loss of macroscopic reversibility, even though melting curves confirmed that the hybridization of individual motifs remained reversible. We hypothesize that ligation freezes the recently described interconnected topology of the hydrogel^27^: individual motifs may dissociate upon heating, but covalently linked loops likely remain topologically entangled, preventing reassembly upon cooling. This frozen architecture also appears to resist salt-dependent denaturation, which could be advantageous when low-salt conditions are required. Future work could explore whether such fixed topologies can be exploited to build imposed structures.

Several of the changes induced by ligation are particularly relevant for biological applications. First, while non-ligated hydrogels progressively ejected captured cells to the gel surface through strand rearrangement, ligation ensured cells remained within the hydrogels.

The suppression of rearrangement ability also translated to changes in material properties: ligated hydrogels no longer exhibited fluorescence recovery, fusion, or relaxation and instead hardened into robust, solid-like materials. By tuning the ligation ratio, we achieved post-formation control over these mechanical properties. This tunability, combined with the enhanced robustness of the ligated gels, facilitates their manipulation and handling. Together, these features make ligated DNA hydrogels attractive candidates for tissue engineering, where precise control over scaffold stiffness is essential to recapitulate different physiological extracellular matrices. In line with this, our 3D-printing experiments suggest that these hydrogels can be shaped into defined architectures suitable for 3D cell culture, complementing prior work on hybrid DNA hydrogels ^33–35^. Although a full rheological characterization would be required to quantitatively establish how ligation regulates the viscoelastic behavior of the gels, such measurements were beyond the scope of this study, which focused on user-oriented properties relevant for manipulation and biological applications. Nevertheless, future work could address this gap and also explore how this hardening could serve as a model for condensate aging, a process implicated in disease-related transitions in cellular biomolecular condensates ^36^.

The stiffness acquired through ligation may also underlie the resistance to DNase I by preventing the distortion of the DNA helices required for the nuclease binding^37^. This is a key advantage, as DNA and RNA cleavage by nucleases plays a critical role in many biological processes ^38^ and the ubiquitous presence of nucleases in physiological environments constitutes a major barrier for the use of DNA structures in biological applications. Strategies encompassing design, nuclease inactivation through medium preheating, or chemical modifications have been developed to address that issue^39^. Structural rigidity is particularly relevant: tetrahedral structures decay about fifty-fold slower that their linear more flexible counterparts in presence of 10% fetal bovine serum, a difference further enhanced by ligation of nicks^40^. Similarly, DNA prisms display a higher resistance to nucleases than their component strands, and ligation keep them intact for more than 8 days^41^. Besides ligation, crosslinking of thymidines using ultraviolet light allowed to lengthen the lifespan of DNA assemblies from 10 to 60 minutes in physiological levels of DNase I^42^. Here, in presence of actin, ligated DNA hydrogels remain stable for at least two weeks in serum-supplemented medium. Interestingly, although actin was previously shown to prevent DNase I digestion of a hybrid hydrogel^43^, the DNase I inhibitor alone was not sufficient to protect non-ligated pure DNA hydrogels, underscoring the protective role of ligation. In the absence of actin, ligated hydrogels undergo progressive digestion starting around day 4, a feature that may be advantageous for applications such as localized drug delivery ^18,44^.

Finally, we showed that ligated DNA hydrogels can capture and maintain viable cells for at least one week, overcoming key limitations of earlier nanostar-based designs. In non-ligated hydrogels, long-term culture was hindered both by the intrinsic rearrangement of the material, which progressively displaced captured cells toward the gel surface (Fig. S6A), and by rapid degradation in nuclease-rich culture media. By suppressing strand rearrangement and enhancing nuclease resistance, ligation enables sustained three-dimensional cell confinement.

This stability creates opportunities to exploit the full DNA nanotechnology toolbox to functionalize or program the gels. As an illustration of this modularity, we functionalized the ligated gels with the RGD peptide, a common integrin-binding motif. Although we did not assess its biological effects, the incorporation of RGD illustrates how ligated hydrogels can readily host additional biochemical functionalities. While ligation suppresses some SE-based reactivity, additional programmability remains possible, for example via enzyme-cleavable sequences or strand-displacement modules that could reverse ligation or trigger dynamic responses.

## MATERIALS AND METHODS

### Products

The sequences of the Y strands were taken from a previously published design^23^ and synthesized commercially. Unmodified and 5’-phosphorylated DNA strands were purchased unpurified from IDT.

Chemically-modified strands (atto488, atto647N or FAM internal modifications) were obtained from Biomers with HPLC purification. Evagreen dye was delivered by Biotium, and T4 Ligase (M0202), ExoIII (M0206) and DNase I-XT (M0570) were purchased from New England Biolabs. DNA hydrogels were formed in non-binding 96-wells plates (Greiner, 655906). Bovine Serum albumin (BSA) was purchased from Sigma-Aldrich (A9418) and Actin protein from Cytoskeleton (AD99).

### DNA hydrogels formation and ligation

Hydrogels were formed isothermally by mixing the three non-phosphorylated Y-strands and their phosphorylated counterparts (total concentration of each Y-strand pair: 5 µM) in PBS supplemented with 5 mg/mL BSA. Solutions were incubated for 1 h at 37 °C under shaking (450 rpm) in non-binding 96-well plates (final volume: 100µl) or, for some experiments (Fig. 2A and B), in microcentrifuge tubes (final volume: 50µl).

Small DNA droplets (Fig. 4C to G) were formed overnight without mixing in honeycomb 96-well plates (Somar Corporation, mw126p096w). A Y2 strand lacking sticky-end and internally modified with a fluorophore (atto647N for Fig. 7 and atto550 otherwise) was added at a 1:100 ratio for visualization.

Unless stated otherwise, the hydrogels containing phosphorylated strands were treated with T4 ligase for 1h after gel formation.

### Microscopy

All fluorescence images were obtained with a Nikon Ti-2 epifluorescence microscope mounted with a Kinetix camera (Teledyne, 01-KINETIX-M-C).

### Nuclease degradation assay

After formation and ligation, hydrogels were incubated at 37°C for 1h with 2µl of either ExoIII (200 U, Fig. 1) or DNase I-XT (4 U, Fig. 6) and the corresponding reaction buffer at 1x. Hydrogels were imaged before enzyme addition and after 1 h of incubation, and their size evolution was quantified using ImageJ.

### Melting and cooling cycles

Hydrogels were prepared with the same DNA concentrations described above in 50 µl Tris-Hcl 20 mM, 150 mM NaCl, 5 mg/mL BSA, 1x EvaGreen, following either the isothermal formation protocol (37 °C, 450 rpm, 1h, in microcentrifuge tubes) or by slow annealing (90 °C to 20 °C, -1 °C/min). The solutions were then transferred in a CFX96 Touch Real-Time PCR system, equilibrated at 20 °C for 5 min, and subjected to three melting-cooling cycles between 20 and 100 °C, at a ramp of 0.5 °C / 2 min.

### FRAP

Small DNA hydrogels were formed overnight in honeycombs well plate without mixing, with 1:100 of a Y2 strand lacking a sticky-end with an internal FAM modification. After ligation, the hydrogels were transferred to an observation chamber (Biorad Frame-Seal, SLF0201) mounted on a Nikon Ti-2 microscope equipped with a diaphragm (TI2-F-FSC) on a heating plate (Tokai Hit, Tpi-TIZGX) set at 45°C. For each acquisition, the diaphragm was manually adjusted to reveal half a bead, which was bleached by exposing it for 30 sec at full LED power using a 20x objective. Recovery was imaged with a 15x objective at 5% LED power, every 5 seconds. Because LED bleaching is not instantaneous, we quantified recovery using normalized fluorescence curves rather than analytical diffusion models. Background fluorescence was subtracted at each time point, and photobleaching during acquisition was corrected using a factor **k(t)** computed from an unbleached control droplet:

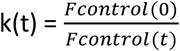

Recovery was computed as:

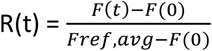

where F_ref, avg_ the mean fluorescence of the unbleached half over the last five time points.

### Fusion assay

DNA droplets were formed and ligated as described for FRAP, then transferred to the same observation chamber and maintained at 45°C. Time-lapse imaging was performed to record droplet encounter and fusion events.

### Mechanical resistance assay

After formation and ligation in non-binding 96-well plates, hydrogels were submitted to increasing speeds of formation in an Eppendorf Thermomixer C.

### 3D printing

DNA strands (500µM each) were mixed in PBS supplemented with 5 mg mL⁻¹ BSA, together with T4 ligase and its buffer (total volume 500 µL), and loaded into a 1.5 mL syringe fitted with either a 21G needle or a conical nozzle (Musashi Engineering; inner diameter 0.1 mm). The syringe was mounted on a Tobeca extrusion printer equipped with a motorized x–y–z stage. Printing was carried out on coverslips treated with poly-lysine, at room temperature and 70% humidity, with an extrusion rate of 0.1 mL h⁻¹ and a working distance of 300 µm, following predefined g-code patterns. The stage translated at 6 cm min⁻¹. After printing, PBS was added to prevent drying of the hydrogel.

### Cell capture and culture

A stable GFP-expressing MCF-7 cell line (Applied Biological Materials, T3904) was used for cell capture experiments. Cells were labelled following our previously published protocol. Briefly, cells were first incubated with a biotinylated anti-EpCAM antibody (eBioscience, 13-9326-82), followed by a streptavidin–DNA conjugate hybridized to three short DNA strands presenting sticky-ends. Labelled cells were mixed with the DNA solution during hydrogel formation, allowing capture through sticky-end hybridization. Hydrogels containing cells were maintained in PriGrow III medium supplemented with 10% FBS, at 37°C and 5% CO_2_.

## CRediT AUTHORSHIP CONTRIBUTION STATEMENT

**Audrey Cochard**: Conceptualization, Data curation, Investigation, Methodology, Project Administration, Supervision, Validation, Visualization, Writing – original Draft, Writing – review and editing. **Lucas Pinède**: Investigation. **Yannick Tauran**: Investigation, Writing – review and editing. **Arnaud Brioude**: Resources. **Beomjoon Kim**: Resources. **Alexandre Baccouche**: Conceptualization, Supervision, Writing – review and editing. **Vincent Salles**: Investigation. **Anthony Genot**: Conceptualization, Funding acquisition, Methodology, Project administration, Resources, Supervision. **Soo Hyeon Kim**: Conceptualization, Funding acquisition, Project administration, Resources, Supervision, Writing – review and editing.

## DATA AVAILABILITY

Data will be made available on request.

## ACKNOWLEDGMENTS

This work was partially supported by Canon Medical Systems Corporation. A.C. is grateful for the support from the CNRS grant PEPR MoleculArXiv (ANR-22-PEXM-0002). The IRP CNRS SupraDNA partially supported Y.T., V.S. and A. Brioude.

---

**Figure S1:**
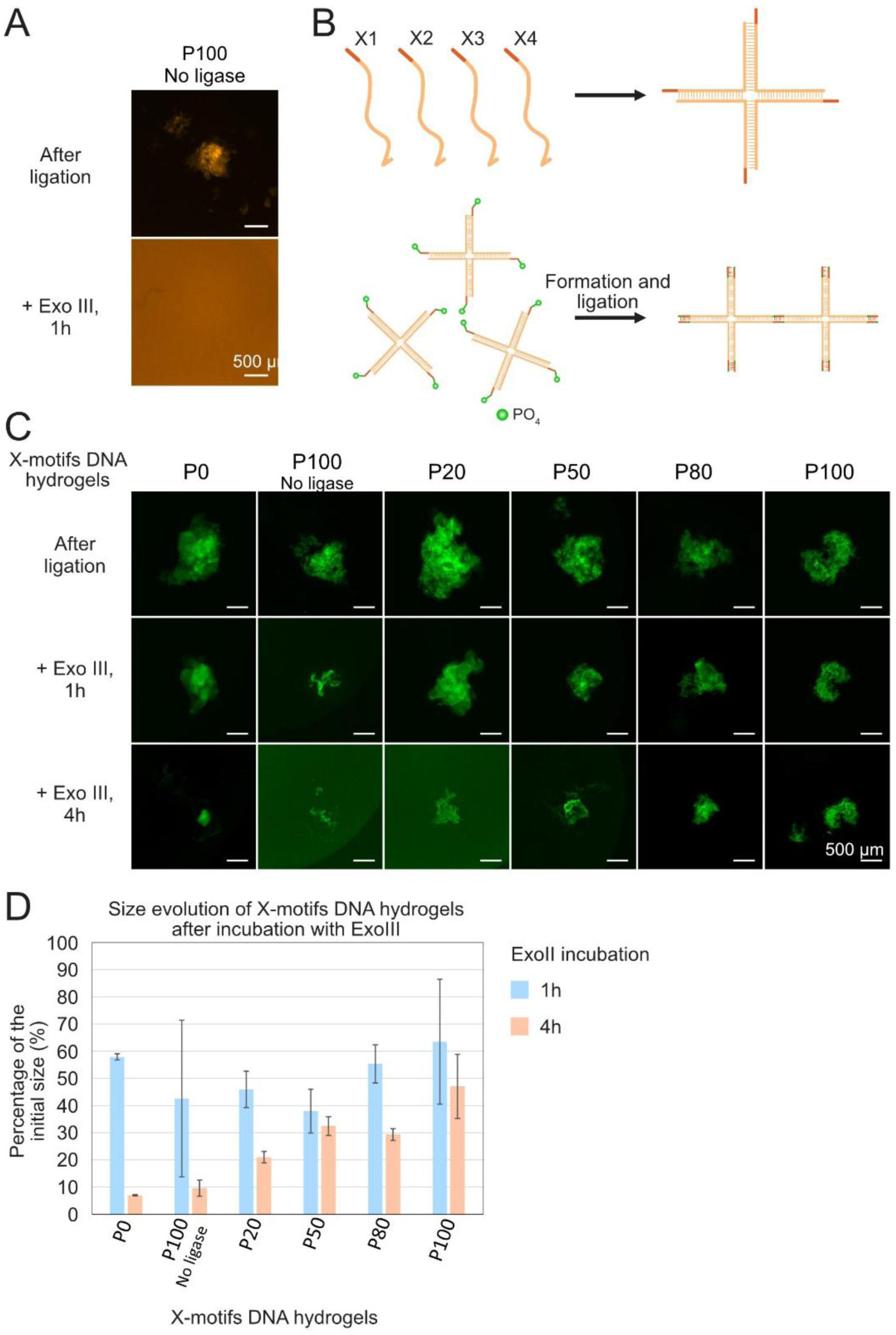
Exonuclease on a non-ligated control hydrogel and formation and ligation of X-motifs DNA hydrogels. **A.** Response of a non-ligated P100 gel to 1h treatment with ExoIII exonuclease. **B.** Schematic of the formation of a DNA nanostar hydrogel based on X-motifs through liquid-liquid phase separation. Here, four complementary DNA strands form X-motifs that become the building blocks of the DNA network. **C.** Response of X-motifs DNA hydrogels with different ratios of phosphorylated strands, after ligation, to 1h and 4h treatment with ExoIII exonuclease. **D.** Size evolution of X-motifs DNA hydrogels following 1h and 4h treatment with ExoIII. N = 2.

**Figure S2:**
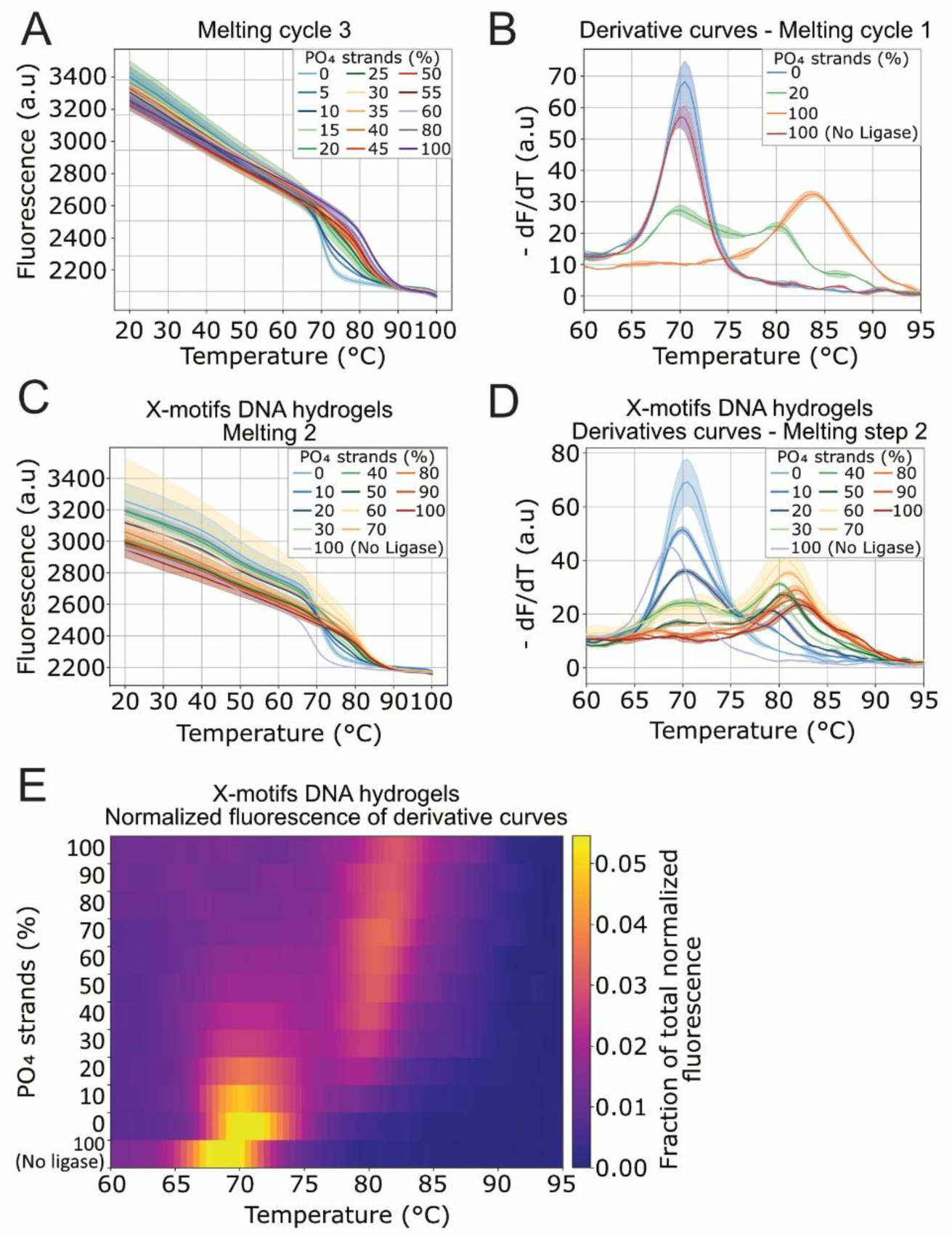
Reversibility of motifs hybridization. **A.** Melting curves of the 3^rd^ cycle for Y-motifs DNA hydrogels (P0 to P100). **B.** Derivatives curves between 60 and 95°C of the 1^st^ melting for Y-motifs DNA hydrogels (P0, P20, P100 and non-ligated P100) **C.** Melting curves of the 2^nd^ cycle for X-motifs DNA hydrogels (P0 to P100). **D.** Derivative curves between 60 and 95°C of the melting curves of the 2^nd^ cycle for X-motifs DNA hydrogels displayed in C. **E.** Normalized derivative fluorescence intensity of the curves displayed in D. For each condition, derivative curves were baseline-corrected and normalized such that the total area under the curve equals 1. The color therefore represents the fractional contribution of each temperature point to the overall derivative fluorescence signal of that condition.

**Figure S3:**
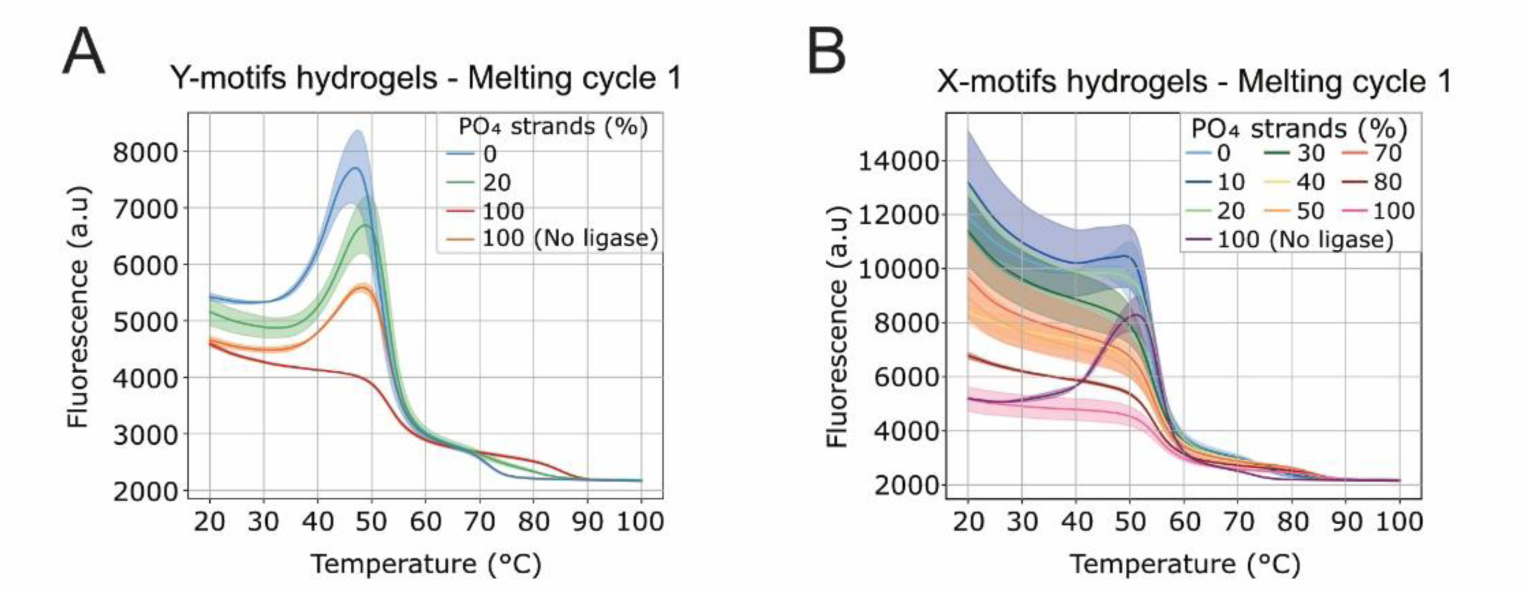
1^st^ melting curves – Dependance on the formation method. **A.** 1^st^ melting curves of Y-motifs DNA hydrogels (P0, P20, P100 and non-ligated P100 gels). **B.** 1^st^ melting curves of X-motifs DNA hydrogels.

**Figure S4:**
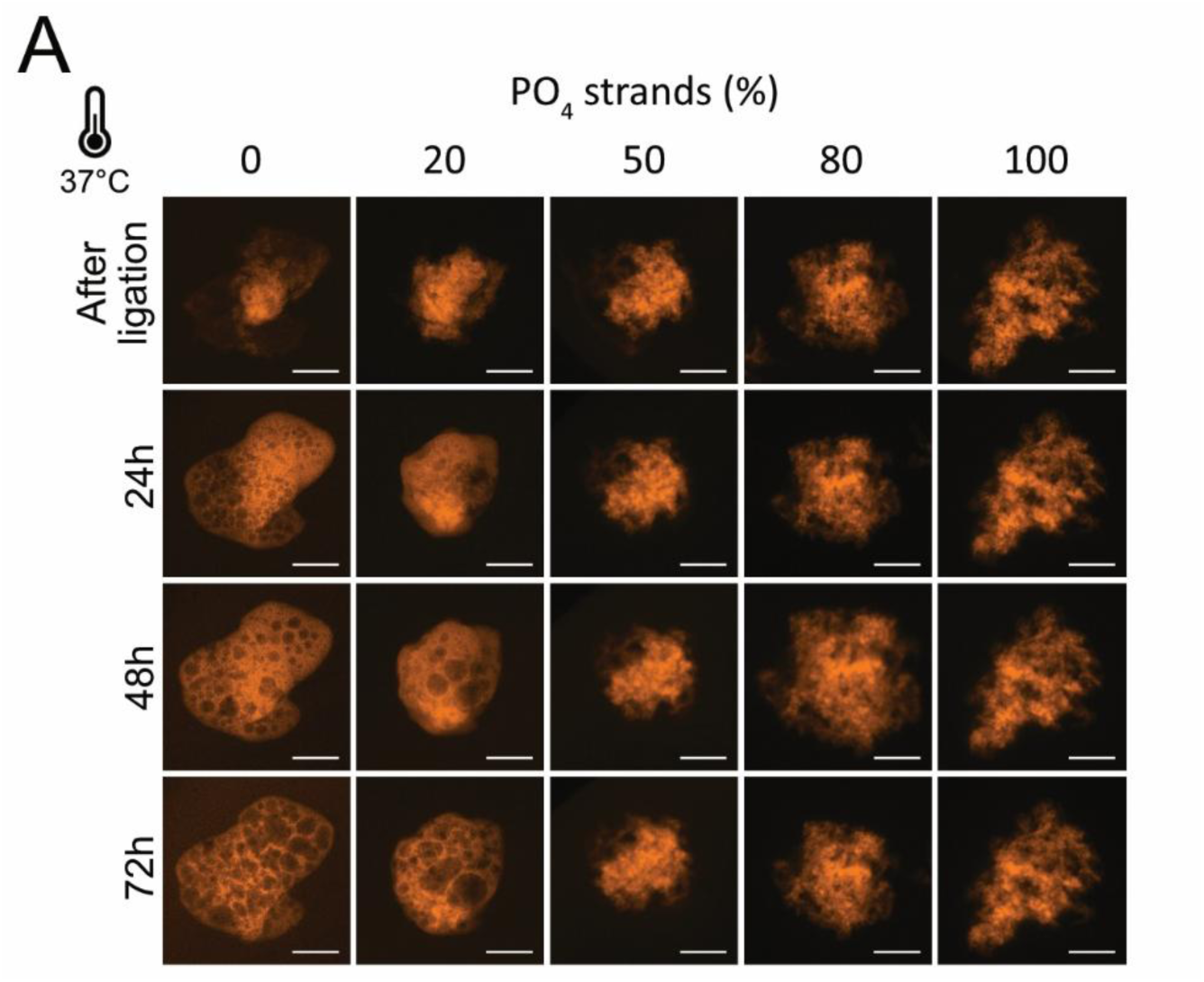
DNA hydrogels rearrangement at 37°C. **A.** Evolution at 37°C for 72h of the DNA hydrogels of Fig. 2A with different percentages of phosphorylated strands, after ligation. Some images were rotated to better highlight the absence of DNA hydrogels shape evolution over time.

**Figure S5:**
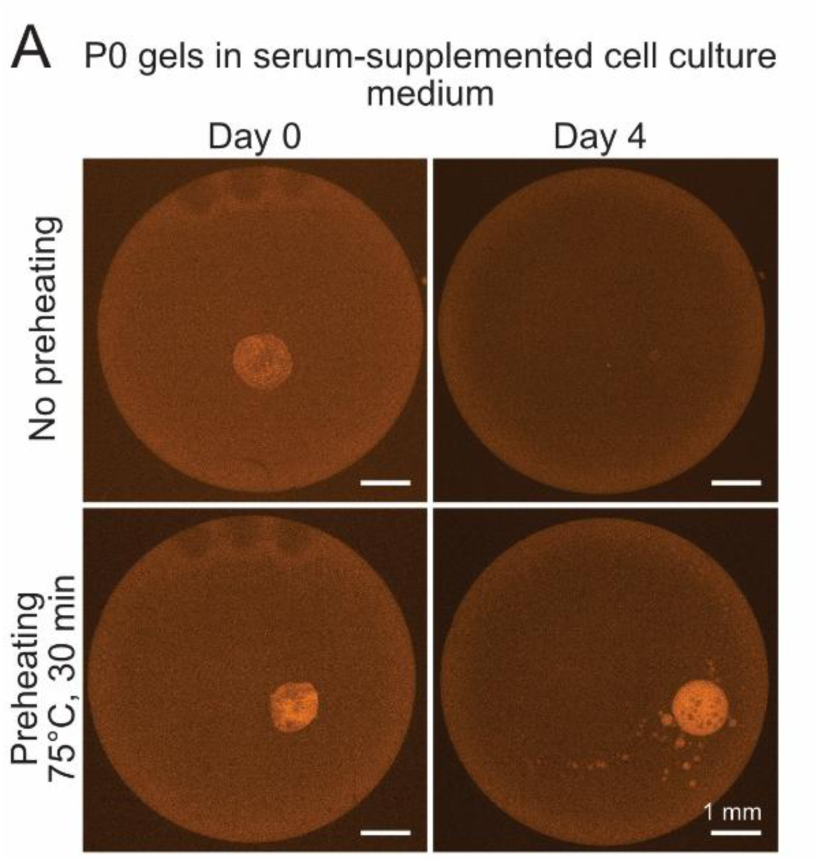
Effect of medium preheating on nuclease resistance. **A.** Evolution over 4 days of P0 gels in serum-supplemented medium either not preheated (top) or preheated (bottom) at 75°C for 30 min.

**Figure S6:**
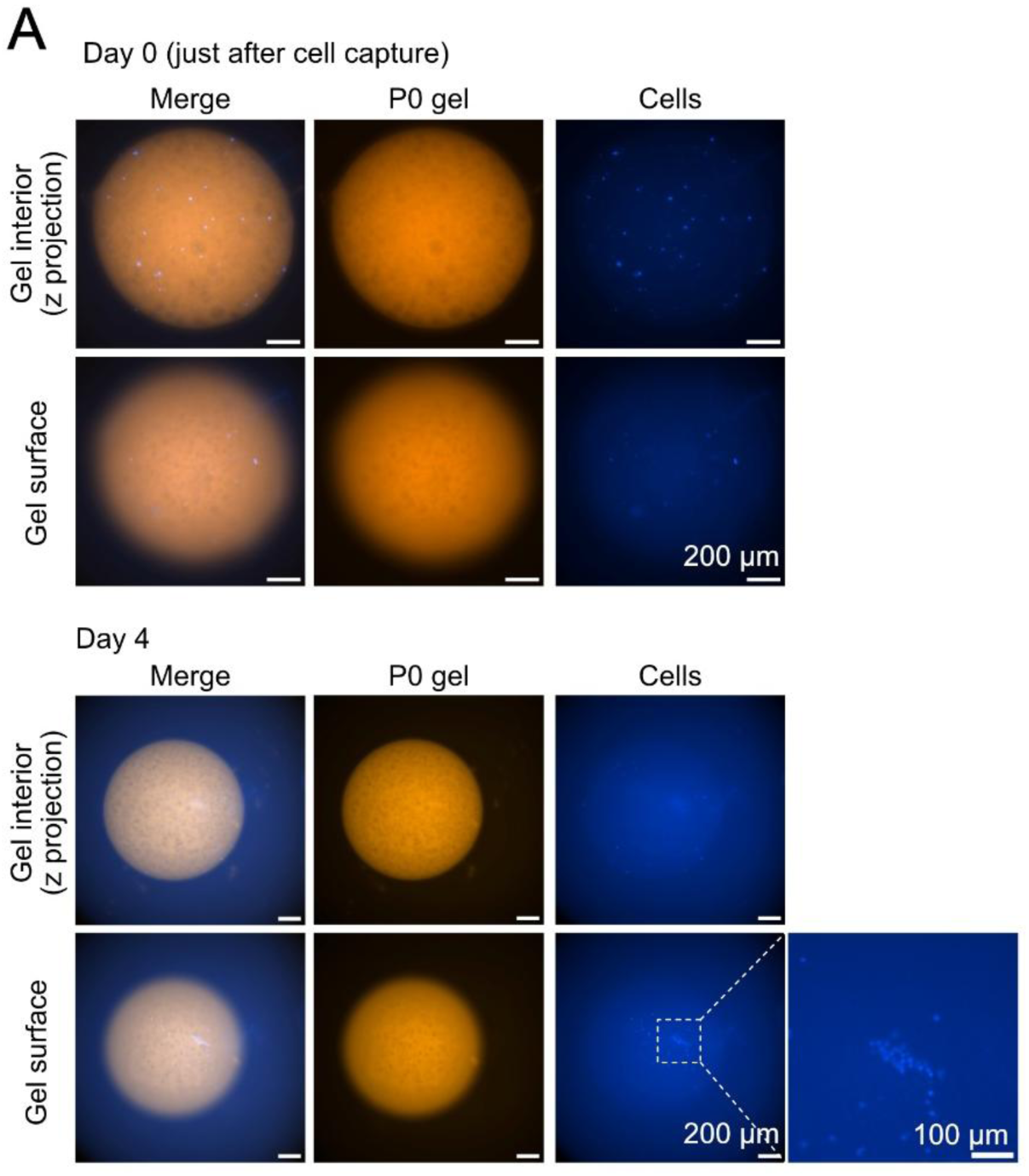
Expulsion of captured cells out of a P0 gel through strand rearrangement. **A.** Fluorescence images of Capan-2 cells (blue, labeled with CMAC Cell Tracker) captured in a P0 gel (orange) right after cell capture (top) and after four days in PBS (down). Images of the gel interior are a projection of z-stack acquisition. The image on the bottom right is a zoom in of the cells at the surface of the gel after 4 days.

